# The D-mannose/L-galactose pathway plays a predominant role in ascorbate biosynthesis in the liverwort *Marchantia polymorpha* but is not regulated by light and oxidative stress

**DOI:** 10.1101/2023.08.16.553627

**Authors:** Tetsuya Ishida, Yasuhiro Tanaka, Takanori Maruta, Takahiro Ishikawa

**Author notes:** **Corresponding author**: Takahiro Ishikawa.

## Abstract

Ascorbate plays an indispensable role in plants, functioning as both an antioxidant and a cellular redox buffer. It is widely acknowledged that the ascorbate biosynthesis in the photosynthetic tissues of land plants is governed by light-mediated regulation of the D-mannose/L-galactose (D-Man/L-Gal) pathway. At the core of this light-dependent regulation lies the *VTC2* gene, encoding the rate-limiting enzyme GDP-L-Gal phosphorylase. The *VTC2* expression is regulated by signals *via* the photosynthetic electron transport system. In this study, we directed our attention to the liverwort *Marchantia polymorpha*, representing one of the basal land plants, enabling us to conduct an in-depth analysis of its ascorbate biosynthesis. The *M. polymorpha* genome harbors a solitary gene for each enzyme involved in the D-Man/L-Gal pathway, including *VTC2*, along with three lactonase orthologs, which may be involved in the alternative ascorbate biosynthesis pathway. Through supplementation experiments with potential precursors, we observed that only L-Gal exhibited effectiveness in ascorbate biosynthesis. Furthermore, the generation of *VTC2*-deficient mutants through genome editing unveiled the inability of thallus regeneration in the absence of L-Gal supplementation, thereby revealing the importance of the D-Man/L-Gal pathway in ascorbate biosynthesis within *M. polymorpha*. Interestingly, gene expression analyses unveiled a distinct characteristic of *M. polymorpha*, where none of the genes associated with the D-Man/L-Gal pathway, including *VTC2*, showed upregulation in response to light, unlike other known land plants. This study sheds light on the exceptional nature of *M. polymorpha* as a land plant that has evolved distinctive mechanisms concerning ascorbate biosynthesis and its regulation.

## INTRODUCTION

Plants harbor elevated concentrations of ascorbate (hereafter referred to as ASC and DHA for the reduced and oxidized forms, respectively), which participate in diverse physiological aspects, functioning not only as an antioxidant but also as a cofactor for various enzymes and an intracellular redox buffer (Smirnoff, 2000; 2018). Within land plants and green algae, the principal mechanism for ascorbate production involves the D-mannose/L-galactose (D-Man/L-Gal) pathway. This pathway comprises eight sequential reactions facilitated by phosphomannose isomerase (PMI), phosphomannomutase (PMM), GDP-D-Man pyrophosphorylase (GMP), GDP-D-Man-3′,5′-epimerase (GME), GDP-L-Gal phosphorylase (GGP), L-Gal-1-P phosphatase (GPP), L-Gal dehydrogenase (GDH), and L-galactono-1,4-lactone dehydrogenase (GLDH) (Figure 1). All the enzymes, except for the final enzyme, GLDH, which localizes in the inner mitochondrial membrane, are distributed within the cytosol. Although certain cytosolic enzymes seem to target both the cytosol and the nucleus, the final precursor L-galactono-1,4-lactone is synthesized in the cytosol and transported to the intermembrane space of the mitochondria, where it undergoes conversion to ascorbate. In contrast to plants, *Euglena gracilis*, a unicellular phytoflagellated protozoan, lacks the D-Man/L-Gal pathway and instead synthesizes ascorbate through a pathway involving D-galacturonic acid (D-GalUA) as a representative intermediate. Aldono-lactonase (ALase) assumes a pivotal role in this pathway, and the *Euglena* ALase has been identified as an orthologue of gluconolactonase (SMP30) in the animal pathway for ascorbate biosynthesis (Ishikawa et al., 2008) (Figure 1). Several studies have postulated a potential role for the D-GalUA pathway in higher plants (Agius et al., 2003; Badejo et al., 2012): however, the lactonase gene remains unidentified, and no conclusive molecular genetic evidence supporting the existence of the D-GalUA pathway has been presented thus far. In contrast to higher plants, the moss *Physcomitrella patens* has two *ALase* genes, but it has been demonstrated that these genes do not actively participate in ascorbate biosynthesis in the moss (Sodeyama et al., 2022). The D-Man/L-Gal pathway prevails as the principal pathway in *P. patens*, as observed in known land plants.

**Figure 1.**
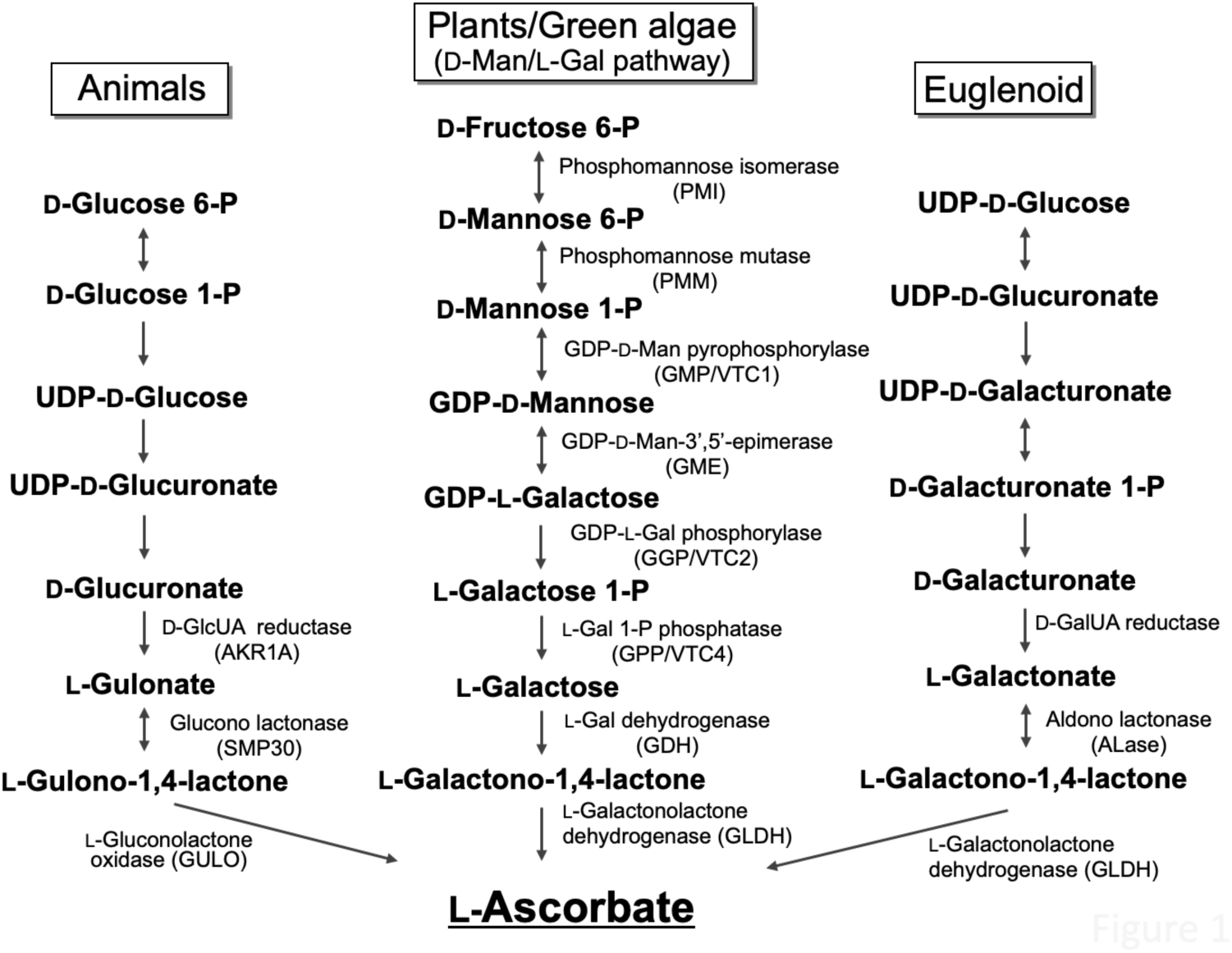
Distribution of major pathways involved in ascorbate biosynthesis across various organisms. The illustration depicting the distribution of ascorbate biosynthesis pathways is based on a previous publication (Wheeler et al., 2015).

Light constitutes the paramount environmental stimulus in the regulation of ascorbate biosynthesis in terrestrial plants. Leaf ascorbate levels escalate in response to light intensity and display diurnal fluctuations, augmenting during daylight hours and diminishing during nocturnal periods. This observation underscores the vital role of a light-responsive regulatory mechanism in ascorbate biosynthesis (Bartoli et al., 2006; Dowdle et al., 2007; Yabuta et al., 2007). Among the enzymes involved in the D-Man/L-Gal pathway, GGP assumes preeminence in the light-mediated control of ascorbate biosynthesis. This enzyme is encoded by two genes, namely *Vitamin C Defective 2* (*VTC2*) and *VTC5*, within the Arabidopsis genome. The transcriptional upregulation of *VTC2* and the augmentation of GGP activity in the light, particularly under high-light conditions, leads to an expansion of the ascorbate content. The light-induced accumulation of ascorbate is predominantly, albeit not completely, impaired in *vtc2* mutants, characterized by a diminished ascorbate content ranging from 20% to 30% of that observed in the wild-type plants (Dowdle et al., 2007; Gao et al., 2011; Terai et al., 2020). A photosynthetic electron transport inhibitor known as 3-(3,4-dichlorophenyl)−1,1-dimethylurea (DCMU) comprehensively suppresses the transcriptional activation of *VTC2* in response to light, suggesting the involvement of an unidentified retrograde signal derived from photosynthesis in the regulation of *VTC2* expression (Yabuta, et al., 2007; Gao et al., 2011). Besides VTC2, VTC3 emerges as a significant component implicated in the light-mediated control of ascorbate biosynthesis. Although the *VTC3* gene does not encode a constitutive enzyme associated with the D-Man/L-Gal pathway, it does encode a distinctive protein harboring both protein kinase and phosphatase domains (Conklin et al., 2000; 2013). The *vtc3* mutant exhibits an approximate 50% reduction in leaf ascorbate content compared to that of wild-type plants, and the illumination does not influence the ascorbate content in the mutant (Conklin et al., 2000; 2013).

Although it has been established that the D-Man/L-Gal pathway operates universally in the biosynthesis of ascorbate in higher plants, knowledge regarding Bryophytes and algae remains limited. Our recent findings demonstrate that the D-Man/L-Gal pathway represents a prevailing pathway and is subject to regulation by light in a photosynthesis-dependent manner in the moss *Physcomitrella patens* (Sodeyama et al., 2021). This implies that light- and photosynthesis-dependent regulation of ascorbate biosynthesis is a common occurrence in terrestrial plants. Conversely, it has been reported that the biosynthesis of ascorbate in the unicellular green alga *Chlamydomonas reinhardii* is not influenced by photosynthesis, despite the presence of the D-Man/L-Gal pathway (Vidal-Meireles et al., 2017). The expression of the *VTC2* gene in *C. reinhardii* is up-regulated in response to external oxidative stress treatment and high-light exposure, leading to an augmentation in ascorbate content (Vidal-Meireles et al., 2017). Thus, it is likely that reactive oxygen species (ROS) act as mediators in activating the biosynthesis of ascorbate in green algae. These observations suggest that the regulation of ASC biosynthesis, which is contingent on light and photosynthesis, evolved in land plants after the acquisition of the D-Man/L-Gal pathway.

It is, therefore, highly intriguing to ascertain how terrestrial plants have evolutionarily acquired the light- and photosynthesis-dependent regulation of ascorbate biosynthesis. In order to obtain further elucidation regarding the evolution of ascorbate biosynthesis regulation, we have directed our attention towards the liverwort *Marchantia polymorpha*, which holds the evolutionary position as the most basal among land plants and has recently emerged as an exceptional model system exhibiting minimal genomic redundancy across its regulatory pathways (Kohchi et al, 2021). In this study, we have successfully demonstrated the conservation and predominant functionality of the D-Man/L-Gal pathway in the liverwort. Interestingly, our observations reveal that the expression of the *VTC2* gene in *M. polymorpha* remains unaffected by light/photosynthesis or oxidative stress, thereby not manifesting any discernible influence on ascorbate biosynthesis.

## RESULTS

### The genome of *Marchantia polymorpha* harbors genes encoding a repertoire of constitutive enzymes in the D-Man/L-Gal pathway and putative lactonases

In order to ascertain the genetic composition of enzymes involved in the process of ascorbate biosynthesis in *M. polymorpha*, a homology search was conducted against MarpolBase (Kawamura et al., 2022; https://marchantia.info/). The BLASTp search successfully identified orthologs that encode a series of enzymes responsible for the D-Man/L-Gal pathway within the liverwort genome (Table 1). It is worth noting that within the *M. polymorpha* genome, all eight enzymes were encoded by individual genes, whereas in other plant species such as Arabidopsis and the moss *Physcomitrium patens* certain enzymes are encoded by two or three paralogous genes (Dowdle et al., 2007; Sodeyama et al., 2021). An intriguing observation arises from the fact that the *M. polymorpha* database did not yield any hits for the *VTC3* ortholog, which encodes a unique enzyme comprised of protein kinase and phosphatase domains. Furthermore, our investigation revealed the presence of three potential genes encoding lactonases, which play a role in ascorbate biosynthesis in both animals and *Euglena gracilis*. The lactonases found in *M. polymorpha* exhibited an approximate 27% identical to rat SMP30 (lactonase) (Table 1). These results suggested that *M. polymorpha* possesses two distinct pathways for ascorbate biosynthesis, namely the D-Man/L-Gal pathway and the D-GalUA pathway.

**Table 1.**
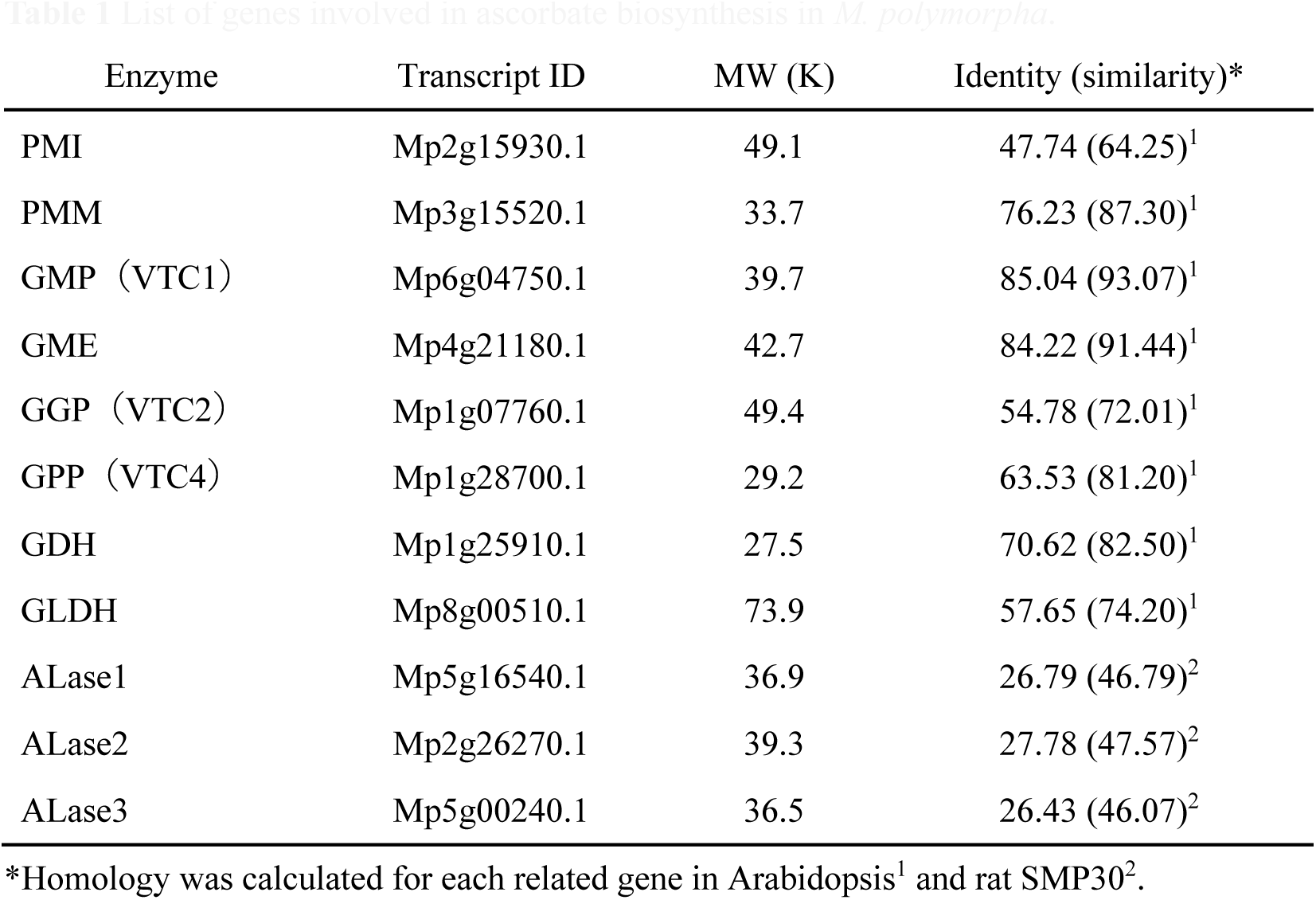
List of genes involved in ascorbate biosynthesis in *M. polymorpha*.

| Enzyme | Transcript ID | MW (K) | Identity (similarity)* |
| --- | --- | --- | --- |
| PMI | Mp2g15930.1 | 49.1 | 47.74 (64.25)<sup>1</sup> |
| PMM | Mp3g15520.1 | 33.7 | 76.23 (87.30)<sup>1</sup> |
| GMP (VTC1) | Mp6g04750.1 | 39.7 | 85.04 (93.07)<sup>1</sup> |
| GME | Mp4g21180.1 | 42.7 | 84.22 (91.44)<sup>1</sup> |
| GGP (VTC2) | Mp1g07760.1 | 49.4 | 54.78 (72.01)<sup>1</sup> |
| GPP (VTC4) | Mp1g28700.1 | 29.2 | 63.53 (81.20)<sup>1</sup> |
| GDH | Mp1g25910.1 | 27.5 | 70.62 (82.50)<sup>1</sup> |
| GLDH | Mp8g00510.1 | 73.9 | 57.65 (74.20)<sup>1</sup> |
| ALase1 | Mp5g16540.1 | 36.9 | 26.79 (46.79)<sup>2</sup> |
| ALase2 | Mp2g26270.1 | 39.3 | 27.78 (47.57)<sup>2</sup> |
| ALase3 | Mp5g00240.1 | 36.5 | 26.43 (46.07)<sup>2</sup> |

\*Homology was calculated for each related gene in Arabidopsis<sup>1</sup> and rat SMP30<sup>2</sup>.

To assess the substantial pathway involved in ascorbate biosynthesis in *M. polymorpha*, we examined the impact of different precursor treatments on ascorbate levels. Truncated thalli at the age of four weeks were supplemented with L-Gal, D-GalUA, or D-GlcUA, each at a concentration of 5 mM. Subsequently, the ascorbate contents were measured over a period of 12 hours under exposure of 50 μmol/m^2^/s of light. Consequently, the application of L-Gal treatment led to a substantial threefold increase in thallus ascorbate content within 12 hours. Conversely, the other compounds displayed no discernible effect (Figure 2). These results indicated that the D-Man/L-Gal pathway operates preferentially over the alternative pathways utilizing D-GalUA or D-GlcUA as precursors.

**Figure 2.**
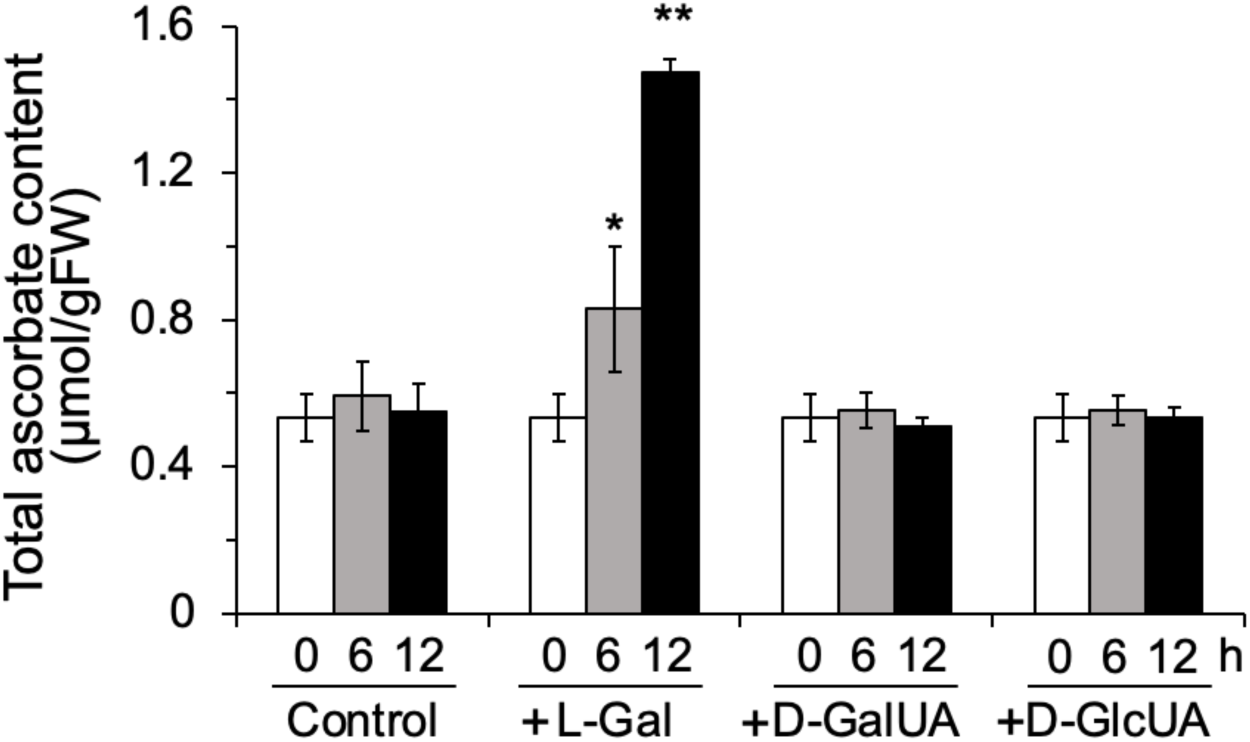
Impact of precursors on the ascorbate content of *M. polymorpha*. Four-week-old thalli were immersed in 5 mM concentrations of L-Gal, D-GalUA, or D-GlcUA and incubated under light conditions at 50 μmol photons m^−2^ s^−1^ for the specified duration. The data are presented as the mean ± SE of three independent experiments. \**P* < 0.05, *\*\*P* <0.01 (vs Control).

### The D-Man/L-Gal pathway represents the predominant route for ascorbate biosynthesis in *M. polymorpha*

To furnish additional compelling substantiation for the function of the D-Man/L-Gal pathway, we endeavored to engender *M. polymorpha* mutants devoid of the *VTC2* gene by CRISPR/Cas9-mediated genome editing. Subsequently, we procured two distinct progeny strains bearing the potential to elicit knockout mutations, namely *vtc2-KO*#3 and *vtc2-KO*#7. Notably, *vtc2-KO*#3 exhibited one nucleotide deletion leading to frameshift mutation near the catalytic domain known as the HIT motif, culminating in premature termination codons (Figure 3A). In the case of *vtc2-KO*#7, the HIT motif was abolished due to a frameshift caused by a single nucleotide insertion (Figure 3A). Significantly, both *vtc2* mutants necessitated L-Gal supplementation to sustain their proliferation, as their growth was stymied in the absence of L-Gal. Restoration of mutant growth was unattainable even with the provision of D-GalUA and D-GlcUA (Figure 3B), which is congruent with the influence not exhibited on ascorbate levels (Figure 2). GDP-L-Gal phosphorylase acts as a catalyst in the conversion of GDP-L-Gal to L-Gal, contingent upon the presence of inorganic phosphate (Pi). Albeit commercial availability of GDP-L-Gal is non-existent, GGP can employ GDP-D-glucose as an alternative substrate. Recent findings have demonstrated that gauging the degradation rates of Pi-dependent GDP-D-glucose serves as a viable substitute for assessing GGP activity (Sodeyama et al., 2021). As shown in Figure 3C, the Pi-dependent degradation activity in the two *vtc2-KO* mutants fell below the threshold of detection.

**Figure 3.**
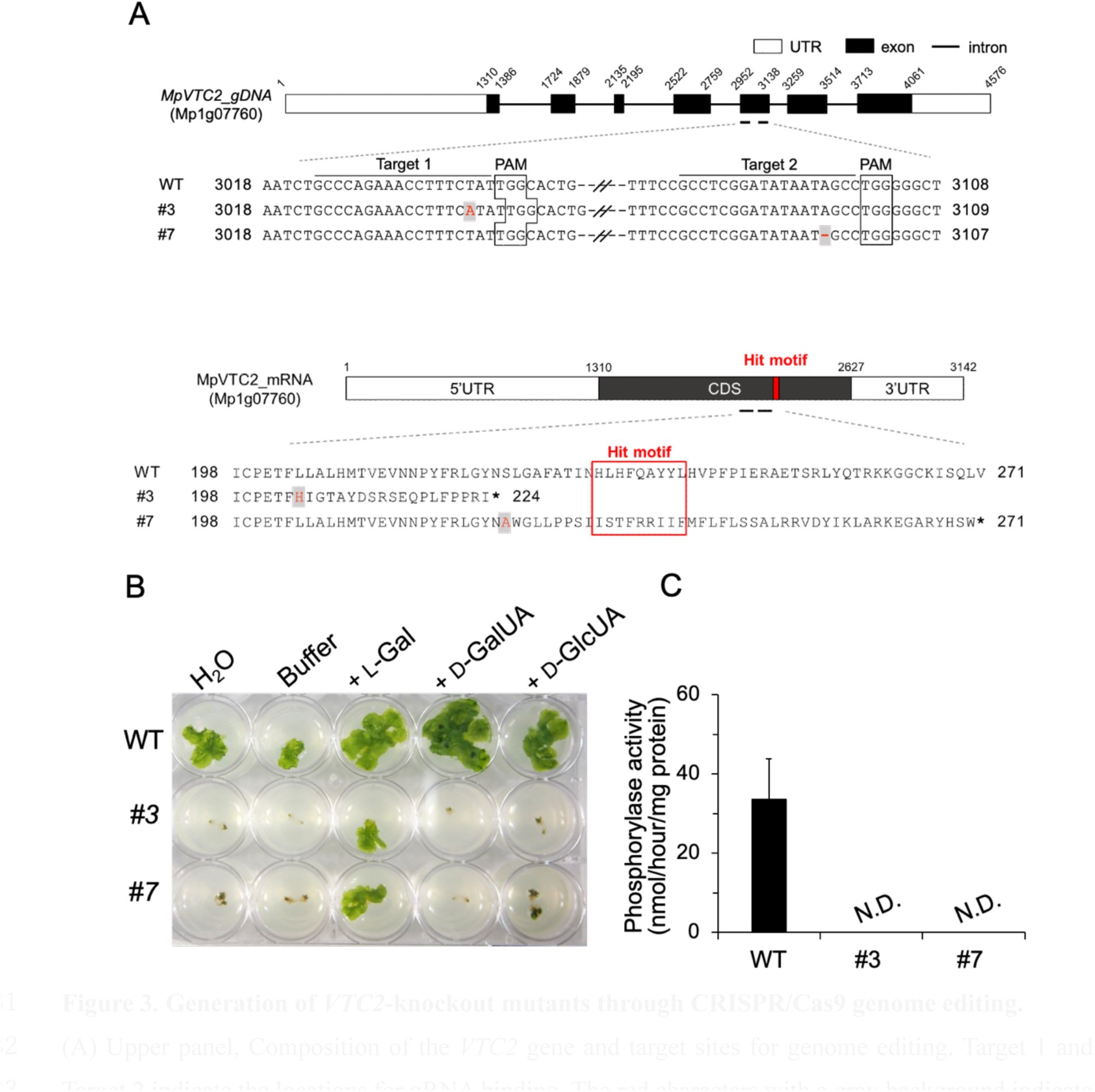
Generation of *VTC2*-knockout mutants through CRISPR/Cas9 genome editing. (A) Upper panel, Composition of the *VTC2* gene and target sites for genome editing. Target 1 and Target 2 indicate the locations for gRNA binding. The red characters with a gray background indicate nucleotide insertion and deletion for lines #3 and #7, respectively. Bottom panel, Overview of *VTC2* mRNA. The Hit motif represents the catalytic domain of GDP-L-Gal phosphorylase. The red characters with a gray background indicate amino acid substitutions resulting from genome editing. (B) Restoration of *VTC2* knockout mutants through supplementation with ascorbate precursors. The panel displays a representative outcome of 4-week-old plants after supplementation with 1 mM of L-Gal, D-GalUA, or D-GlcUA. (C) Evaluation of GDP-L-Gal phosphorylase activity. Phosphorylase activities were measured in crude extracts prepared from 4-week-old thalli of WT and *VTC2* knockout mutants (#3 and #7) using GDP-D-glucose as an alternative substrate. The data are presented as the mean ± SE of three independent experiments. N.D., not detected.

We subsequently investigated the impacts of VTC2 on ascorbate content. The wild-type plants and the two *vtc2-KO* mutants were cultivated for a duration of four weeks in a medium supplemented with 1 mM L-Gal. Subsequently, they were transferred to a medium devoid of L-Gal and were further cultivated under a light regime consisting of a 16-hour light and an 8-hour dark cycle, with a light intensity of 50 μmol/m^2^/s. In the case of the wild type, the levels of ascorbate in the thallus, immediately prior to the transfer to the L-Gal free medium, were recorded to be 0.74 ± 0.02 μmol/gFW. These levels remained relatively constant throughout the experiment after transplantation, at approximately 0.60 ± 0.06 μmol/g FW (Figure 4A). The ascorbate levels in the *vtc2-KO* #3 and #7 were 0.55 ± 0.08 and 0.52 ± 0.04 μmol/g FW, respectively, in the presence of the L-Gal supplemented medium. These levels were found to be lower than those observed in the wild type. Notably, in contrast to the wild type, the ascorbate contents in the KO mutants exhibited a significant decrease, reaching 0.23 ± 0.03 μmol/g FW (approximately 40% of the wild-type value) by 8 days after transplantation onto L-Gal free medium. This decline was followed by a gradual decrease, reaching 0.10 ± 0.01 μmol/g FW by 36 days (Figure 4A). These findings provide compelling genetic evidence supporting the dominant role of the D-Man/L-Gal pathway in the liverwort *M. polymorpha*.

**Figure 4.**
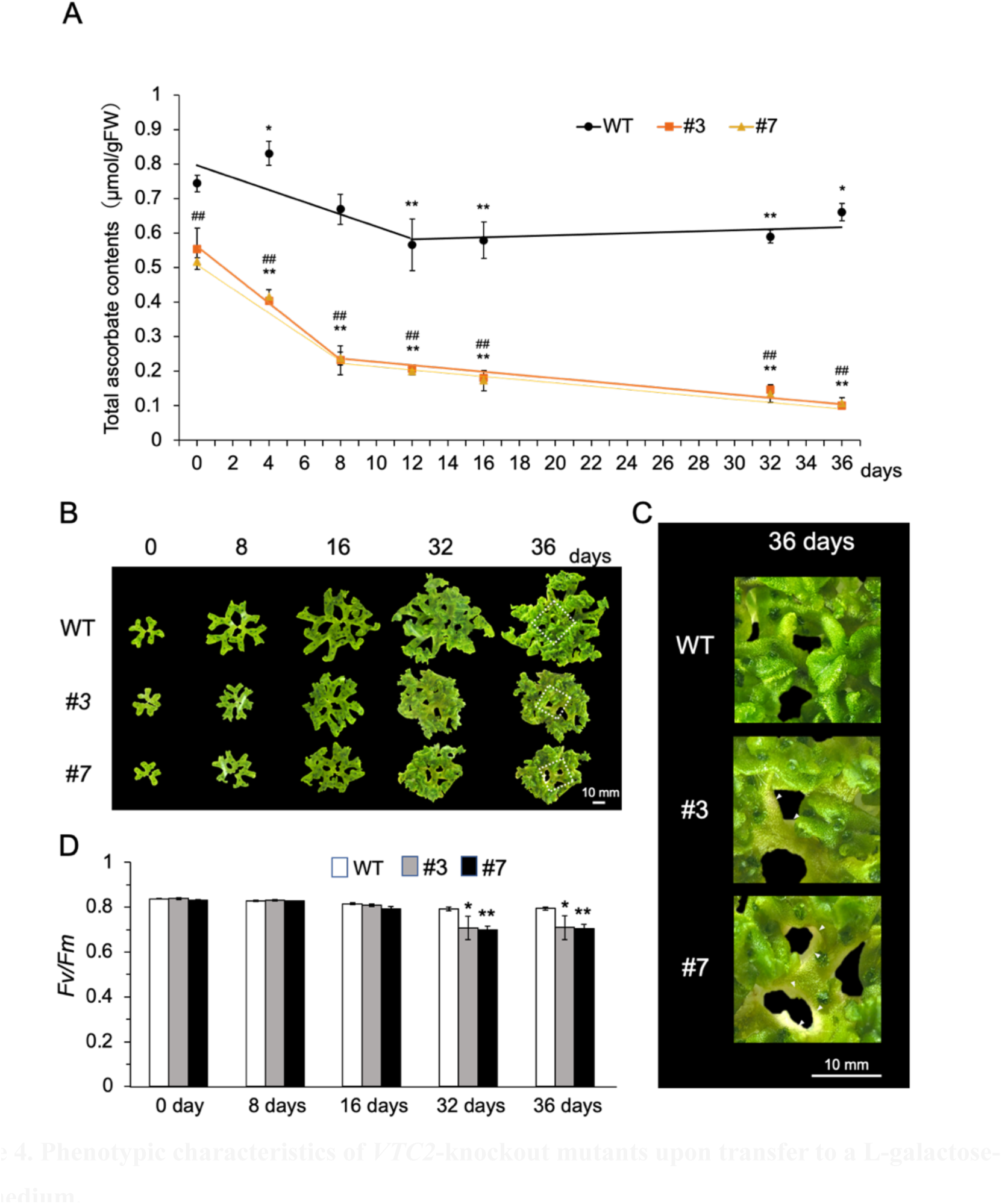
Phenotypic characteristics of *VTC2*-knockout mutants upon transfer to a L-galactose-free medium. Four-week-old plants cultivated in a medium supplemented with 1 mM L-Gal were transferred to a L-Gal-free medium and maintained under standard growth conditions with an light intensity of 50 μmol photons m^−2^ s^−1^. Measurements of each parameter were taken on the specified days. (A) Alterations in ascorbate content of thalli. The data are presented as the mean ± SE of three independent experiments. \**P* < 0.05, *\*\*P* <0.01 (compared to day 0); #*p*<0.05, *##p*<0.01 (compared to WT) (B) Visible phenotype of *VTC2*-knockout mutants (#3 and #7). (C) Enlarged view of the white dashed enclosure of the day 36 plants in Panel B. (D) The PSII activity (Fv/Fm) was determined at 22°C after dark adaptation for 30 min. The data are presented as the mean ± SE of three independent experiments. \**P* < 0.05, *\*\*P* <0.01 (compared to WT).

Upon transferring to an L-Gal-free medium, the magnitude of thallus in the KO mutants consistently exhibited smaller dimensions compared to the wild type. Moreover, this disparity tended to escalate over time. Notably, by the 16th-day post-transplantation, no discernible contrast emerged in the maximum quantum yield of Photosystem II (Fv/Fm) values, a representative marker of oxidative stress, despite a gradual decline in ascorbate concentrations up to this juncture (Figure 4B). However, from the 32nd day onward, a substantial decline in Fv/Fm became apparent in the KO mutant, concomitant with the onset of chlorosis at the thallus center (Figures 4C and 4D). These findings compellingly underscore the indispensability of ascorbate biosynthesis for robust thallus growth. Based on these results, we have inferred that a minimum of 0.2 µmol/g FW of ascorbate is imperative to avert photoinhibition.

### Transcription of genes involved in the D-Man/L-Gal pathway remains uninduced in *M. polymorpha* upon exposure to light

In higher plants and the moss *Physcomitrella patens*, the ascorbate concentration in tissues engaged in photosynthesis increasees in response to light intensity. This phenomenon aligns with the transcriptional up-regulation of some genes encoding enzymes pertaining to the D-Man/L-Gal pathway, especially *VTC2*, which is contingent upon photosynthetic electron transport system (Dowdle et al., 2007; Yabuta et al., 2007; Yoshimura et al, 2014; Sodeyama et al., 2021). Consequently, we investigated the effect of light on the biosynthesis of ascorbate in *M. polymorpha*. First, the ascorbate content and the expression levels of genes responsible for D-Man/L-Gal pathway enzymes were evaluated under normal growth light conditions (50 µmol photons m^−2^ s^−1^). The initial content of total ascorbate prior to exposure to light was measured as 0.40 ± 0.04 µmol/gFW (Figure 5A), a markedly inferior value when compared to the documented ranges in Arabidopsis and *P. patens*, which fall between 1 and 5 µmol/gFW (Shiroma et al., 2019; Sodeyama et al., 2021). Additionally, the proportion of DHA to total ascorbate in *M. polymorpha* amounted to approximately 60%, significantly surpassing the corresponding values observed in higher plants and *P. patens* (approximately 10-20%). Thus, the ascorbate redox status in this plant is considerably low. Following 3 hours of exposure to light, the cumulative content of total ascorbate escalated to 0.52 ± 0.02 µmol/gFW, approximately 1.3 times greater than the pre-illumination measurement (Figure 5A). Although a statistically significant distinction was observed between the pre- and post-illumination values, the disparity remained inconsequential in comparison to Arabidopsis and *P. patens*.

**Figure 5.**
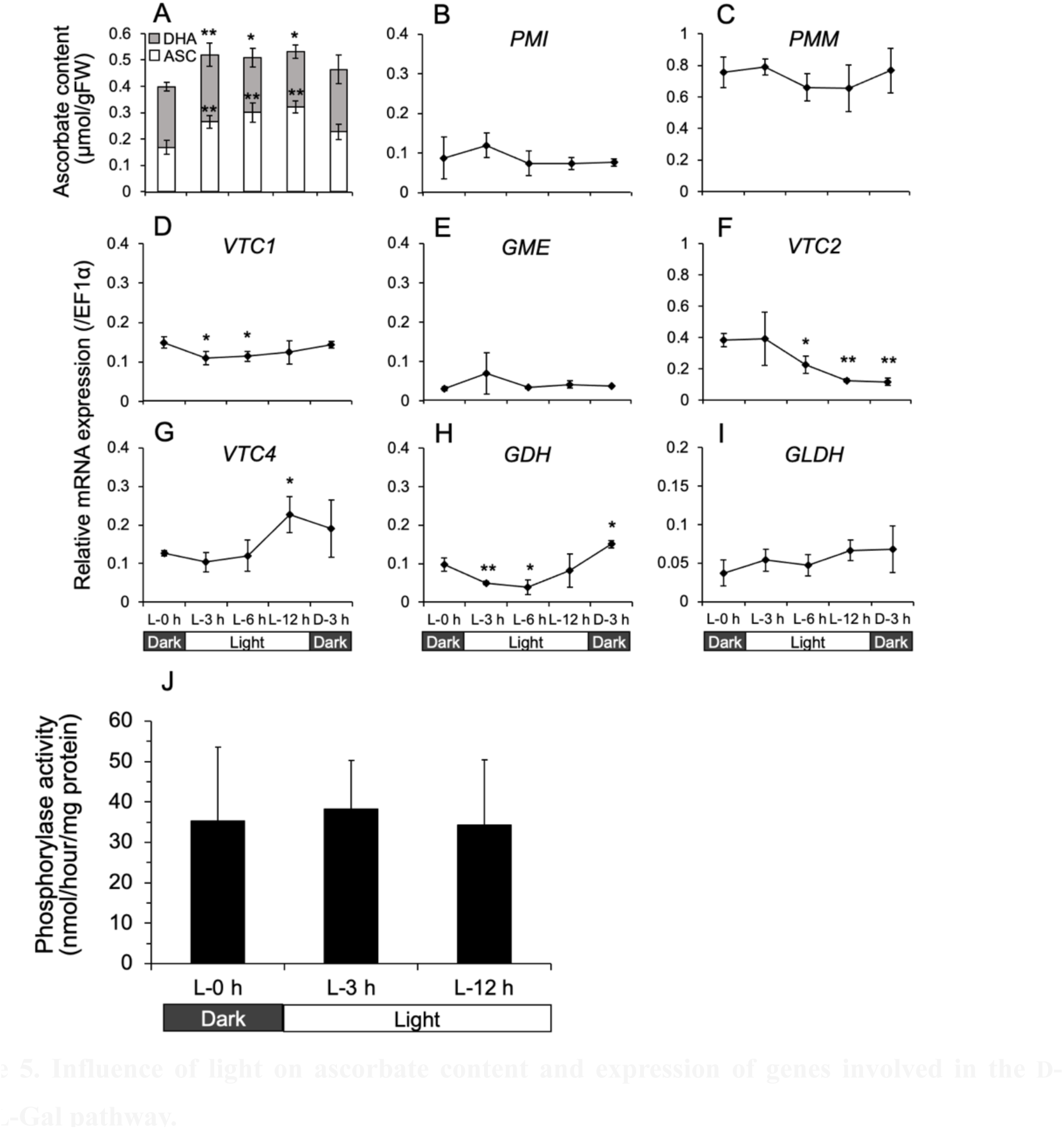
Influence of light on ascorbate content and expression of genes involved in the D-Man/L-Gal pathway. Four-week-old *M. polymorpha* plants grown on plates were subjected to a 12h light/12h dark cycle with a light intensity of 50 μmol photons m^−2^ s^−1^. Sampling was conducted at each indicated time point, and the levels of ascorbate and the expression of each gene were assessed. The data are presented as the mean ± SE of three independent experiments. (A) Ascorbate content. (B)-(I) Expression of genes related to the D-Man/L-Gal pathway. (J) Assessment of GDP-L-Gal phosphorylase activity. Phosphorylase activities were quantified in crude extracts derived from 4-week-old thalli using GDP-D-glucose as an alternative substrate. The data are presented as the mean ± SE of three independent experiments. \**P* < 0.05, *\*\*P* <0.01 (compared to L-0h).

None of the genes exhibited any prominent alterations in expression three hours after exposure to light, despite a slight augmentation in ascorbate levels (Figures 5B-I). Among the biosynthetic genes, the transcriptional activity of the *VTC4* gene, responsible for encoding GPP, was the sole gene that manifested a noteworthy increase after 12 hours of light exposure. However, this increment did not coincide temporally with the surge in ascorbate levels observed at 3 hours (Figure 5G). Instead, transcript levels of various genes (*VTC1*, *VTC2*, and *GDH*) were significantly reduced following light irradiation (Figures 5D, 5F, and 5H). Furthermore, the activity of GGP remained constant and exhibited no variation after light irradiation (Figure 5J). In higher plants and *P. patens*, DCMU treatment, which inhibits the photosynthetic electron transport chain, effectively suppresses the light-induced accumulation of ascorbate and the transcriptional activity of *VTC2* (Yabuta et al., 2007; Sodeyama et al., 2021). In *M. polymorpha*, DCMU treatment curtails the light-induced buildup of ascorbate while leaving the transcription of *VTC2* unaffected, thus suggesting that the suppression of ascorbate accumulation can be attributed to the depletion of sugar (resulting from photosynthesis inhibition), which serves as the substrate for its biosynthesis (Figure S1).

We conducted additional experiments to examine the effect of prolonged periods of light exposure at varying intensities on ascorbate levels and the transcription of the *VTC2* gene. To accomplish this, we subjected four-week-old *M. polymorpha* plants to continuous light at intensities of either 50 or 600 µmol photons m^−2^ s^−1^ (referred to as GL or HL, respectively) for a duration of 48 hours. Comparative analysis revealed that HL irradiation had a more pronounced effect on increasing ascorbate levels when compared to GL (Figure S2). However, neither GL nor HL demonstrated a significant influence on *VTC2* gene transcription (Figure S2). These results indicate that genes associated with the D-Man/L-Gal pathway in *M. polymorpha* are not governed by light and photosynthesis.

### The D-Man/L-Gal pathway in *M. polymorpha* is not influenced by oxidative stress

Vidal-Meireles et al., (2017) have reported that in the green alga *Chlamydomonas reinhardtii*, which also biosynthesizes ascorbate through the D-Man/L-Gal pathway, the regulation of ascorbate biosynthesis is independent of light and photosynthesis. Instead, they found that the expression of the enzymes involved in the D-Man/L-Gal pathway, including *VTC2*/GGP, is significantly augmented under conditions of oxidative stress, such as treatments with H_2_O_2_ or Rose Bengal, the latter of which functions as a singlet oxygen generator. These observations prompted our investigation into the effect of oxidative stress on ascorbate biosynthesis in *M. polymorpha*. Four-week-old liverwort plants derived from asexual shoots were subjected to treatments with 1 mM and 10 mM H_2_O_2_ to assess changes in ascorbate content and *VTC2* gene expression levels over time. Concerning the ascorbate content, it tended to decrease at 3 hours following the treatment with 1 mM H_2_O_2_ compared to the control, and no subsequent increase was noted thereafter (Figure 6A). Under the 10 mM H_2_O_2_ treatment, the ascorbate content exhibited a significant decrease to approximately 40% at 3 hours post-treatment and continued to decline thereafter (Figure 6A). The *VTC2* gene expression levels were assessed by qPCR at the 3-hour time point for each H_2_O_2_ concentration. Consequently, these treatments failed to induce an elevation in the transcript levels of the *VTC2* gene (Figure 6B). Additionally, comparable results were obtained with the administration of Rose Bengal (Figure S3). These results unequivocally demonstrate that oxidative stress does not activate *VTC2* gene expression in *M. polymorpha*.

**Figure 6.**
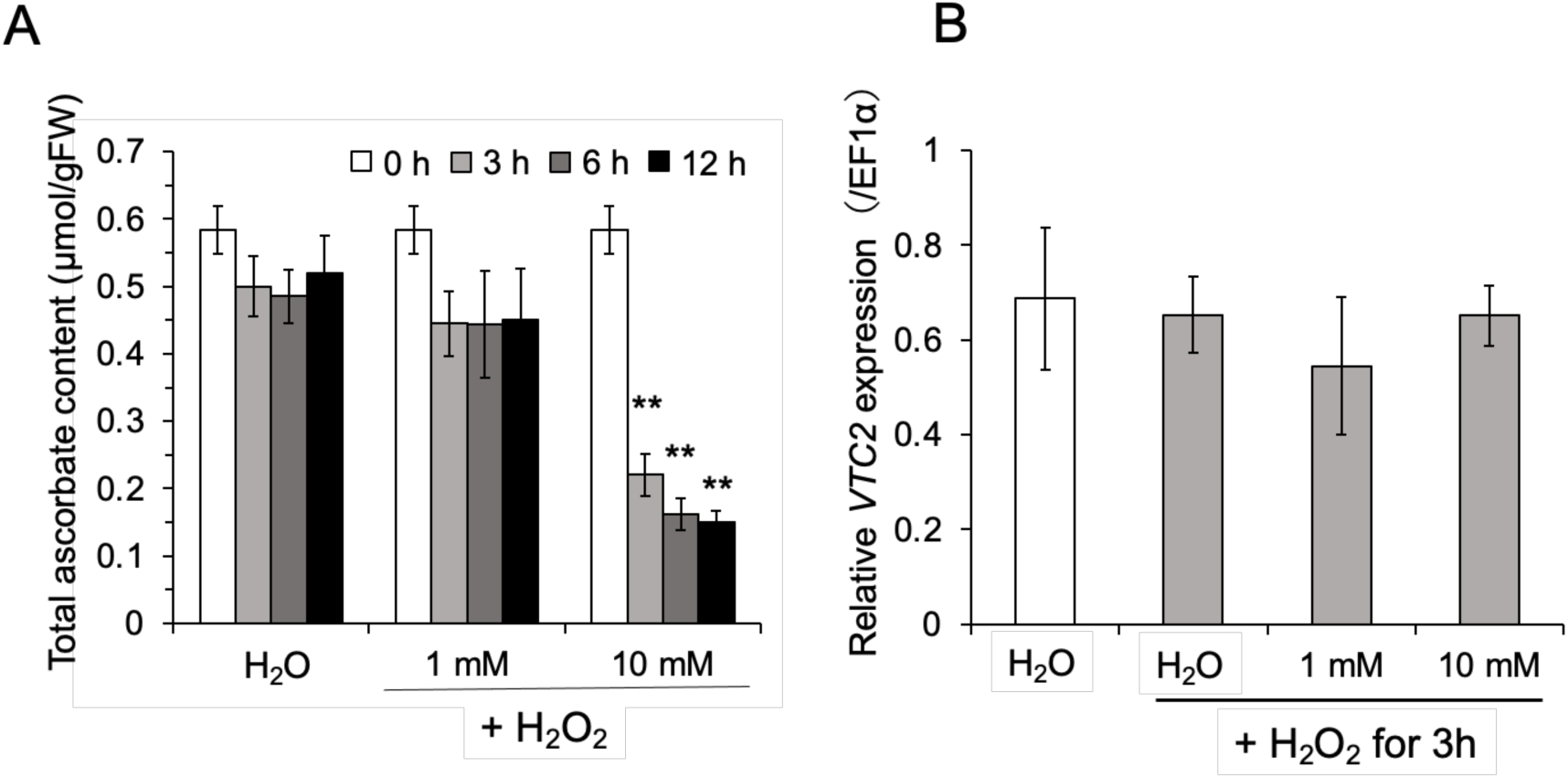
Impact of H_2_O_2_ treatment on ascorbate content and *VTC2* gene expression. (A) Ascorbate content. Four-week-old *M. polymorpha* plants grown on plates were treated with 1 mM or 10 mM H_2_O_2_ at a light intensity of 50 μmol photons m^−2^ s^−1^. Sampling was performed at each indicated time point, and the total ascorbate content was measured. The data are presented as the mean ± SE of three independent experiments. \**P* < 0.05, *\*\*P* <0.01 (compared to H_2_O). (B) *VTC2* expression level. Hydrogen peroxide treatment was administered by immersing the plants in 1 mM or 10 mM H_2_O_2_ for 3 h at a light intensity of 50 μmol photons m^−2^ s^−1^. The expression levels of the *VTC2* gene were evaluated using quantitative real-time PCR. The data are presented as the mean ± SE of three independent experiments.

## DISCUSSION

### Ascorbate biosynthesis pathway and its significance in *M. polymorpha*

In plans, apart from the D-Man/L-Gal pathway, three alternative pathways for ascorbate biosynthesis have been postulated, employing D-galacturonate, L-gulose, and *myo*-inositol as representative metabolic intermediates (Smirnoff, 2018; Ishikawa et al., 2018). Among these pathways, only the D-Man/L-Gal pathway has thus far garnered compelling physiological evidence based on molecular genetics, with all its constituent enzymes identified at the genetic level (Smirnoff, 2018; Ishikawa et al., 2018). The D-Man/L-Gal pathway encompasses eight enzymatic steps, and certain enzymes involved in these steps, such as PMI, GMP, and GGP, are encoded by multiple paralogous genes in terrestrial plants. For example, Tao et al., (2020) have reported that most plant species, including gymnosperms, monocotyledons, gymnosperms, bryophytes, and bryophyte lineages, possess at least two *VTC2* homologs, owing to various duplications throughout their evolutionary history. Conversely, *M. polymorpha*, in line with genomic information indicating minimal gene duplication, possesses only one gene for each of the enzymes in the D-Man/L-Gal pathway (Bowman, et al., 2017). Thus, *M. polymorpha* proves advantageous for molecular physiological investigations of the D-Man/L-Gal pathway, as the need to consider functional complementation by paralogs is obviated. In this study, we demonstrated that the absence of a solitary *VTC2* gene establishes the dominance of the D-Man/L-Gal pathway in ascorbate biosynthesis in *M. polymorpha*. The *vtc2* knockout mutants were unable to thrive without exogenous L-Gal supplementation. In the case of Arabidopsis and *P. patens*, which possess two or more genes encoding GGP, a singular loss of *VTC2* does not yield lethal effects. Only when *VTC2* and its paralog (*VTC5*) are simultaneously disrupted, an ascorbate-dependent growth phenotype is observed in Arabidopsis (Dowdle et al., 2007).

Ascorbate plays a pivotal physiological role not only as an antioxidant but also as an intracellular redox buffer and enzyme cofactor (Smirnoff, 2018). As an illustration, ascorbate acts as a coenzyme for 2-oxoglutarate-dependent dioxygenases, which participate in the synthesis of phytohormones like ethylene, gibberellins, and abscisic acids, as well as in the biosynthesis of hydroxyproline for extracellular matrix glycoproteins such as extensins and arabinogalactan-proteins (Smirnoff, 2018). Hence, a complete deficiency of ascorbate can prove fatal to plants during the vegetative developmental stage due to multifaceted factors necessitating ascorbate. Nevertheless, it is noteworthy that moss plants, having reached a certain degree of growth with an adequate supply of ascorbate, can sustain normal growth for approximately one month under conditions where ascorbate is not provided, despite lacking the capacity for ascorbate biosynthesis. This phenomenon is likely facilitated through ascorbate recycling (the reduction of the oxidized form, DHA, to ASC). Actually, *M. polymorpha* has multiple isoform genes for both monodehydroascorbate reductase (MDAR) and DHA reductase (DHAR). Our current data indicate that a minimum of approximately 0.2 µmol/gFW of ascorbate is necessary for the healthy growth of the liverwort under our laboratory growth conditions.

In contrast to the genes involved in the D-Man/L-Gal pathway, we found three paralogous genes encoding lactonase in *M. polymorpha*. This attribute is of interest due to the rarity of gene duplication in the liverwort. This investigation has not yet examined whether these gene products function as lactonases. Furthermore, there is currently no experimental evidence indicating that lactonases contribute to ascorbate biosynthesis in terrestrial plants. However, the role of the enzyme lactonase in ascorbate biosynthesis has been thoroughly elucidated in mice and the microalgae *Euglena gracilis*, wherein ascorbate is synthesized through D-GlcUA and D-GalUA as intermediates, respectively (Kondo et al., 2006; Ishikawa et al., 2008). Among land plants, the moss *P. patens* possesses two lactonase genes, while they have never been identified in any vascular plants. Similar to the *Euglena* enzyme, both *P. patens* lactonases (PpALase 1 and 2) exhibit activities for catalyzing the conversion of L-galactonic acid to the ultimate precursor L-galactono-1,4-lactone. Although the kinetic properties suggest that PpALase1 is physiologically functional, the absence of this enzyme does not result in a decrease in ascorbate content. Instead, in the *PpALase1* knockout mutants, an accumulation of DHA occurs, leading to a cell death phenotype accompanied by a decline in ascorbate redox status. This phenomenon aligns with the fact that PpALase1 possesses lactonase activity for DHA (Sodeyama et al., 2021). Consequently, it has been suggested that PpALase1 may serve a crucial role in DHA degradation rather than ascorbate biosynthesis. The involvement of *M. polymorpha* lactonases in DHA degradation remains unknown, and this possibility contradicts the fact that approximately 60% of the total ascorbate in the liverwort is in the form of DHA. Nevertheless, the presence of lactonases exclusively in bryophytes within land plants is intriguing, warranting further meticulous analysis to ascertain the physiological functions of the three lactonase orthologs in *M. polymorpha*.

### The expression of the *VTC2* gene in *M. polymorpha* exhibits no responsiveness to either light or oxidative stress

In higher plants, GGP represents the rate-limiting step in the D-Man/L-Gal pathway and has garnered significant attention as a target for modification. This includes endeavors to develop vitamin C hyperaccumulator plants as well as abiotic and biotic stress-tolerant plants by enhancing ascorbate content (Bulley et al., 2012; Zhang et al., 2015; Koukounaras, et al., 2022). The activity of GGP is subject to regulation by light at multiple stages, encompassing transcription, translation, and post-translation. Notably, its transcriptional regulation is reliant on photosynthesis. Unraveling these underlying mechanisms will offer valuable insights not only into the fundamental aspects of ascorbate biosynthesis but also its practical applications. Interestingly, in contrast to known land plants, the transcription of the *VTC2* gene in *M. polymorpha* exhibited no response to light and photosynthesis. Furthermore, light did not affect GGP activity, suggesting that GGP in *M. polymorpha* is unlikely to be regulated by light at the translation or post-translational levels. This lack of light responsiveness extended beyond *VTC2* and was also observed in other biosynthetic genes. Nonetheless, total ascorbate levels experienced a slight increase under light conditions, possibly attributed to enhanced substrate availability from photosynthesis to the biosynthetic pathway. Additionally, while *VTC2* transcription is activated by oxidative stress rather than light and photosynthesis in green algae, the *VTC2* gene in *M. polymorpha* did not respond to oxidative stress. These findings indicate that ascorbate biosynthesis in *M. polymorpha* is not governed by either light or oxidative stress.

The transcription of the *VTC2* gene is subject to regulation by oxidative stress in green algae and by light in mosses and higher plants, indicating a transition in the regulatory mechanism of gene expression during plant evolution. In this context, it is intriguing to observe that *VTC2* in *M. polymorpha* exhibits no response to either light or oxidative stress. It is possible that the liverwort has lost the regulatory mechanism of *VTC2* in response to light or oxidative stress throughout its evolutionary history. However, the limited knowledge regarding the *cis*- and *trans*-factors involved in the transcriptional regulation of the *VTC2* gene in land plants and green algae makes it challenging at this time to discuss the absence of transcriptional regulation in the liverwort *VTC2* from this perspective. With regards to the *cis*-element of the *VTC2* gene, Li et al., (2013) have inferred that a G-box motif serves as a light-responsive element through deletion analysis of the *VTC2* promoter in kiwifruits. Since the G-box motif is a ubiquitous *cis*-element and is also present in the promoter region of the liverwort *VTC2* gene, it does not appear to be a critical factor contributing to the deficiency in light responsiveness. Our previous analysis of the Arabidopsis *VTC2* promoter has suggested that a region between −70 to - 40 relative to the transcription start site, which does not contain any known *cis*-elements including the G box, plays a role as a light-responsive *cis*-element (Gao et al., 2011). Therefore, we consider the possibility that the lack of unidentified *cis*-element(s) in the liverwort *VTC2* might contribute to impaired light regulation.

In the case of *M. polymorpha*, it is noteworthy that the transcription of *VTC2* is not stimulated by light, and the levels of ascorbate are not significantly influenced by light. This absence of light responsiveness in ascorbate biosynthesis aligns intriguingly with the fact that *M. polymorpha* lacks the *VTC3* gene. The *VTC3* gene encodes a unique chimeric enzyme containing protein kinase and PP2C-type protein phosphatase domains (Conklin et al., 2013). While this protein exhibits high conservation among land plants and green algae, *M. polymorpha* represents the sole exception to our knowledge. The Arabidopsis *vtc3* mutant exhibits approximately 50% of the ascorbate content compared to the wild type, while the elevation of ascorbate levels in response to light exposure is markedly inhibited (Conklin et al, 2000, 2013). Similar impacts on the ascorbate content due to VTC3 have been observed in *P. patens* mutants with disrupted *VTC3* genes (Sodeyama et al., 2021). Thus, VTC3 seems to play a role in the light-mediated regulation of ascorbate biosynthesis in land plants. The regulatory influence of VTC3 on *VTC2* at the transcriptional and post-translational levels is presently unknown. Exploring whether the absence of *VTC3* is one of the factors influencing light-induced *VTC2* transcription and ascorbate levels in *M. polymorpha* would be of great interest. Furthermore, conducting comprehensive comparisons between *M. polymorpha* and other land plants could facilitate the identification of genes involved in the regulation of *VTC2* expression.

A further notable aspect of ascorbate biosynthesis in *M. polymorpha* resides in its diminutive pool size. The concentration of ascorbate in the illuminated thallus of *M. polymorpha* spans from approximately 0.4 to 0.5 µmol/gFW, which accounts for about one-tenth to one-third of the corresponding levels found in the vascular plant *Arabidopsis thaliana*, where the range lies between 1.5 and 5.0 µmol/gFW (Dowdle et al, 2007). Also, the ascorbate content in *M. polymorpha* amounts to approximately one-sixth to one-half of that contained in the protonemata of the bryophyte *Physcomitrium patens*, measuring approximately 1.0 to 3.0 µmol/gFW (Sodeyama et al., 2021). Through our analysis of the *vtc2-KO* mutants, it has been estimated that *M. polymorpha* necessitates a minimum essential quantity of around 0.2 μmol/gFW of ascorbate for its growth. To the best of our knowledge, *M. polymorpha* possesses the most diminutive ascorbate pool size among all land plants reported thus far. Given its preference for shade and its ability to thrive in humid conditions, it is hypothesized that *M. polymorpha* can withstand such modest levels of ascorbate due to its reduced susceptibility to environmental fluctuations, including light and drought. Additionally, the maintenance of this limited ascorbate pool size may be facilitated by ascorbate recycling systems such as the ascorbate-glutathione cycle. Consequently, *M. polymorpha* could potentially survive without acquiring a light-regulation mechanism for ascorbate biosynthesis.

## CONCLUSION

In this study, we have unequivocally illustrated that the process of ascorbate biosynthesis in the liverwort *M. polymorpha* is orchestrated by the D-Man/L-Gal pathway. This revelation stems from a series of experiments employing strains with disrupted *VTC2* gene and precursor treatments. Notably, our research has also unveiled a distinctive trait of *M. polymorpha*, setting it apart from other terrestrial plants and the green alga *C. reinhardtii*. Unlike its counterparts, the regulation of the *VTC2* gene in *M. polymorpha* is unaffected by light-induced signaling or ROS. Furthermore, the absence of the *VTC3* gene, coupled with the presence of three lactonase orthologs, alludes to the exceptional evolutionary trajectory of *M. polymorpha* with regards to ascorbate metabolism. Our findings posit *M. polymorpha* as a terrestrial model plant of scholarly significance, providing invaluable insights into the evolutionary foundations of ascorbate biosynthesis and its mechanisms of light regulation.

## MATERIALS AND METHODS

### Plant materials and growth conditions

*Marchantia polymorpha* Takaragaike-1 (Tak-1, male accession) was utilized as the wild type specimen. The strain was graciously provided by Prof. Takayuki Kohchi from the Graduate School of Biostudies at Kyoto University, Japan. A half-strength Gamborg’s B5 (1/2×B5) medium was conventionally employed for cultivation. *M. polymorpha* was cultivated on plates containing 1/2×B5 medium solidified with 1.0% agar, maintaining at a temperature of 21°C under a long day condition (16/8 h day/night cycle), and exposed to a light intensity of 50 μmol photons m^−2^ s^−1^.

### Precursor treatment

Four-week-old thallus grown on 1/2×B5 agar medium subjected to treatments using 5 mM concentrations of L-Gal, D-GalUA, and D-GlcUA individually, while 10 mM MES buffer (pH 5.5) served as the negative control. The thalluses were subsequently incubated under illumination at 50 μmol photons m^−2^ s^−1^ and at a temperature of 21°C. The total ascorbic acid level was measured for the collected thalluses collected at 0, 6 and 12 h, respectively.

### Ascorbate measurement

Liverwort samples (approximately 200 mg) were rapidly frozen in liquid nitrogen and ground in 2 mL of 2% (w/v) meta-phosphoric acid. The homogenate was then centrifuged at 13,000 rpm for 20 min at 4°C. The resulting supernatant was transferred to a new tube and immediately utilized for the ASC assay. ASC content was quantified using an ultra-fast liquid chromatography system (Prominence UFLC; Shimadzu, Japan) equipped with a C-18 column (LUNA C18-2, 150×4.6 nm; Shimadzu). The mobile phase comprised 1% meta-phosphoric acid, and the flow rate was set at 0.5 mL min^−1^. ASC was detected at 254 nm using the SPD-20A UV-VIS detector (Shimadzu). Total ascorbate content was measured after the reduction of DHA through incubation with 10 mM of Tris (2-carboxyethyl) phosphine hydrochloride (TCEP; Wako, Japan). The DHA content was calculated by subtracting ASC from the total ascorbate content.

### Inhibitor and stress treatment

Four-week-old thallus grown on 1/2×B5 agar medium underwent treatments with DCMU and oxidative stress. For the DCMU treatment, 3 mL of 0.1% DMSO (Control) or 1, 10, and 100 μM DCMU were administered via spraying onto the thallus after dark adaption. The plants were then subjected to a dark treatment for 1 h and transferred to normal growth conditions (50 μmol photons m^−2^ s^−1^, 21°C). For the high-light treatment, the thallus was exposed to 600 μmol photons m^−2^ s^−1^ for the indicated time after being exposed to normal growth light for 3h. Hydrogen peroxide treatment involved immersing the thallus in 1 mM or 10 mM H_2_O_2_ for the indicated time under the normal growth light.

### GGP activity measurement

A *M. polymorpha* thallus (approximately 400 mg wet/wt) was pulverized in liquid nitrogen and added to 2 mL of 50 mM Tris-HCl (pH 7.5) containing 10 mM MgCl_2_, 2 mM dithiothreitol (DTT), 1 mM aminocapronic acid, 1 mM benzamidine, 1mM phenylmethanesulfluoride (PMSF), and 10% glycerol. The homogenate was centrifuged at 13,000 rpm for 20 min at 4°C. The resulting supernatant was transferred to an ultra-filtration tube (Amicon Ultra0.5 30K; Merck) and centrifuged at 13,000 rpm for 20 min at 4°C to obtain a concentrated enzyme solution for GGP activity measurement. The reaction mixture (100 μL) consisted of 50 mM Tris-HCl (pH 7.5), 2 mM MgCl_2_, 10 mM KCl, 1 mM DTT, 0.1 mM GDP-D-glucose as an alternative substrate for GDP-L-Gal, enzyme (250 μg/mL), and 5 mM sodium phosphate, and was incubated for 30 min at 26°C. The enzyme reaction was terminated by heat treatment for 3 min at 98°C. The resulting reaction mixture was then analyzed using a UFLC system (Prominence; Shimadzu) equipped with a C-18 column (LUNA C18-2, 150×4.6 nm; Shimadzu). The mobile phase consisted of 50 mM Sodium phosphate (pH6.7), 3.5% (v/v) MeOH, 5 mM tetra butyl ammonium hydrogen sulfate, and 5 mg/L EDTA/4Na, with a flow rate of 1.0 mL min^−1^. GDP-D-glucose was detected at 254 nm using the SPD-20A UV-VIS detector (Simadzu). GGP activity was calculated based on the phosphate-dependent decrease in the substrate.

### Plasmid preparation and generation of *M.polymolpha VTC2* knockout mutants

Plasmids for *VTC2* genome editing with CRISPR/Cas9 were constructed following the methodology described by Sugano et al., (2018). MpVTC2-Target1-F and MpVTC2-Target1-R, as well as MpVTC2-Target2-F and MpVTC2-Target2-R, were individually incubated at 95°C for a duration of 5 min each. Subsequently, the heat block was deactivated and allowed to cool to room temperature. The resulting oligonucleotides were then inserted into the pMpGE_En03 plasmid, which had been treated with the *BsaI* restriction enzyme. The accuracy of the sequence was confirmed by DNA sequencing using ABI PRISM 3100xl Genetic Analyzer. The segment between *att*L1 and *att*L2 of the entry vector was subsequently integrated into the binary vector pMpGE010 using the Gateway LR Clonase Enzyme Mix (Thermo Fisher Scientific, USA). The final constructs were introduced into *Agrobacterium tumefaciens* C58 by electroporation. In order to transform *M. polymolpha*, regenerating thalli were co-cultured with the transformed *A. tumefaciens* C58 for 3 days, following the method outlined by Kubota et al., (2013). The transfected thalli were then selected on a 1/2×B5 medium supplemented with 10 μg/mL hygromycin and 100 μg/mL Cefotaxime. The genome sequence of the thalli that survived in the selective medium was examined using DNA sequencing performed with the ABI PRISM 3100xl Genetic Analyzer.

### RNA isolation and cDNA synthesis

Total RNA was extracted from approximately 50 mg of *M. polymorpha*. Initially, the samples were homogenized in liquid nitrogen, and then 1 mL of RNAiso Plus (Takara Bio., Japan) was added and thoroughly mixed. The homogenate was centrifuged at 13,000 rpm for 5 min. Subsequently, 250 μL of chloroform was added to the supernatant, and the samples were well-mixed and centrifuged at 13,000 rpm for 20 min. The resulting supernatant was transferred to a fresh tube, followed by the addition of 1 mL of isopropanol. The mixture was inverted 10 times to ensure proper mixing. After standing for 20 min, the samples were centrifuged at 13,000 rpm for 20 min. The supernatant was discarded, and the pellets were dried using a vacuum dryer (TAITEC, Saitama, Japan). The crude RNA was treated with DNase to eliminate genomic DNA. The concentration and quality of the total RNA were assessed using a NanoDrop Q5000 (Thermo Fisher Scientific). The first strand cDNA was synthesized utilizing reverse transcriptase (ReverTra Ace; Toyobo, Japan) and an oligo dT primer. The reaction was carried out according to the instructions provided by the manufacturer.

### Quantitative real-time PCR

Expression levels of ascorbate biosynthesis genes were quantified through analysis using the LightCycler 96 System (Roche, Switzerland). For quantitative real time PCR, cDNA (20 ng) was combined with 10 μL of 2×GeneAce SYBR^®^ qPCR Mix α No ROX (Nippon gene, Japan) and 1 μL of 1 μM mixed primer (forward and reverse). The total volume was adjusted to 20 μL with H_2_O. Expression levels were normalized using elongation factor 1α (*EF1α*; Mp3g23400) as the reference gene. Please refer to Table S1 for the list of all primers employed. Each experiment was conducted in triplicate, utilizing independently isolated RNA samples.

### Chlorophyll fluorescence measurements

Chlorophyll fluorescence parameters were assessed in thalli following a 30-min period of dark adaptation at 22°C using the FluorCam 800MF (Photon Systems Instruments, Czech Republic). The actinic irradiance was set at 120 μmol photons m^−2^ s^−1^. The *Fv/Fm* ratio and the quantum yields of photosystem II (ΦPSⅡ) were calculated according to the methods described by Yabuta et al., (2007) and Genty et al., (1989).

### Statistical analysis

All experiments were replicated at least three times with three technical replicates. The significance of differences between datasets was evaluated using Student’s t-test. Calculations were performed on data derived from three or more independent biological replicates, utilizing Microsoft Excel.

### Data Availability

All pertinent data can be found within the manuscript and its accompanying supplementary materials.

## ACKNOWLEDGEMENTS

We express our gratitude to Professor Takayuki Kohchi for generously providing *Marchantia polymorpha* and the vectors required for liverwort transformation.

## AUTHOR CONTRIBUTIONS

T. Ishikawa formulated and designed the experiments; T. Ishida, YT and T. Ishikawa conceived the research plans; T. Ishida conducted the majority of the experimental procedures; T. Ishida, YT, TM and T. Ishikawa analyzed and interpreted the data; T. Ishikawa and TM authored the manuscript with contributions from all co-authors. All authors participated in the manuscript editing process. All authors reviewed and approved the final version of the manuscript.

## CPMPETEING INTERESTS

The authors declare that they have no conflict of interest.

## SUPPORTING MATERIALS LEGENDS

Supplementary Table S1. List of primers used for real-time PCR and genotyping.

Supplementary Figure S1. Effect of DCMU treatment on ascorbate content and *VTC2* gene expression in *M.polymorpha*.

Supplementary Figure S2. Effect of high-light exposure on ascorbate content and *VTC2* gene expression in *M.polymorpha*.

Supplementary Figure S3. Effect of photosensitizer Rose Bengal treatment on ascorbate content and *VTC2* gene expression in *M.polymorpha*.

