## Supplementary figures and images for "The D-mannose/L-galactose pathway plays a predominant role in ascorbate biosynthesis in the liverwort *Marchantia polymorpha* but is not regulated by light and oxidative stress"

### Supplemental Figure S1

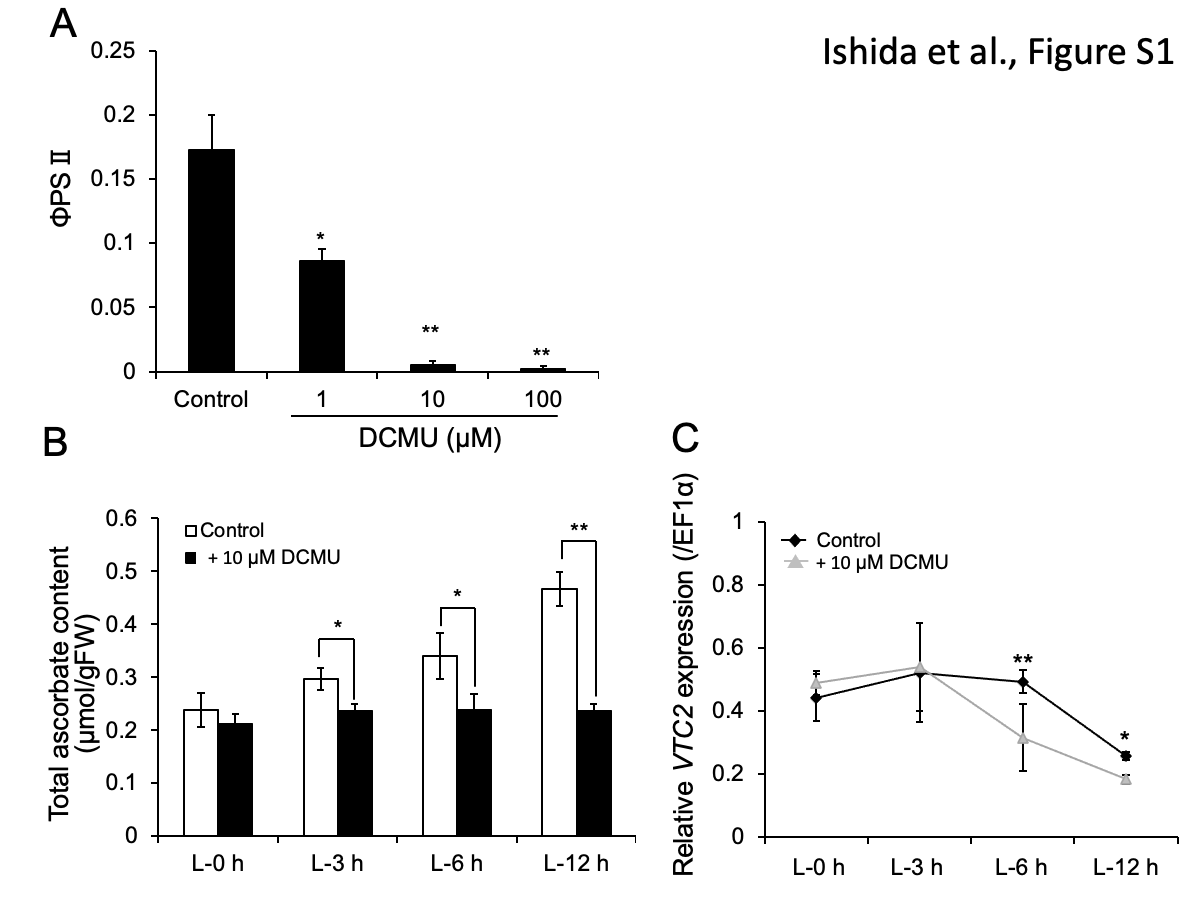

### Supplemental Figure S2

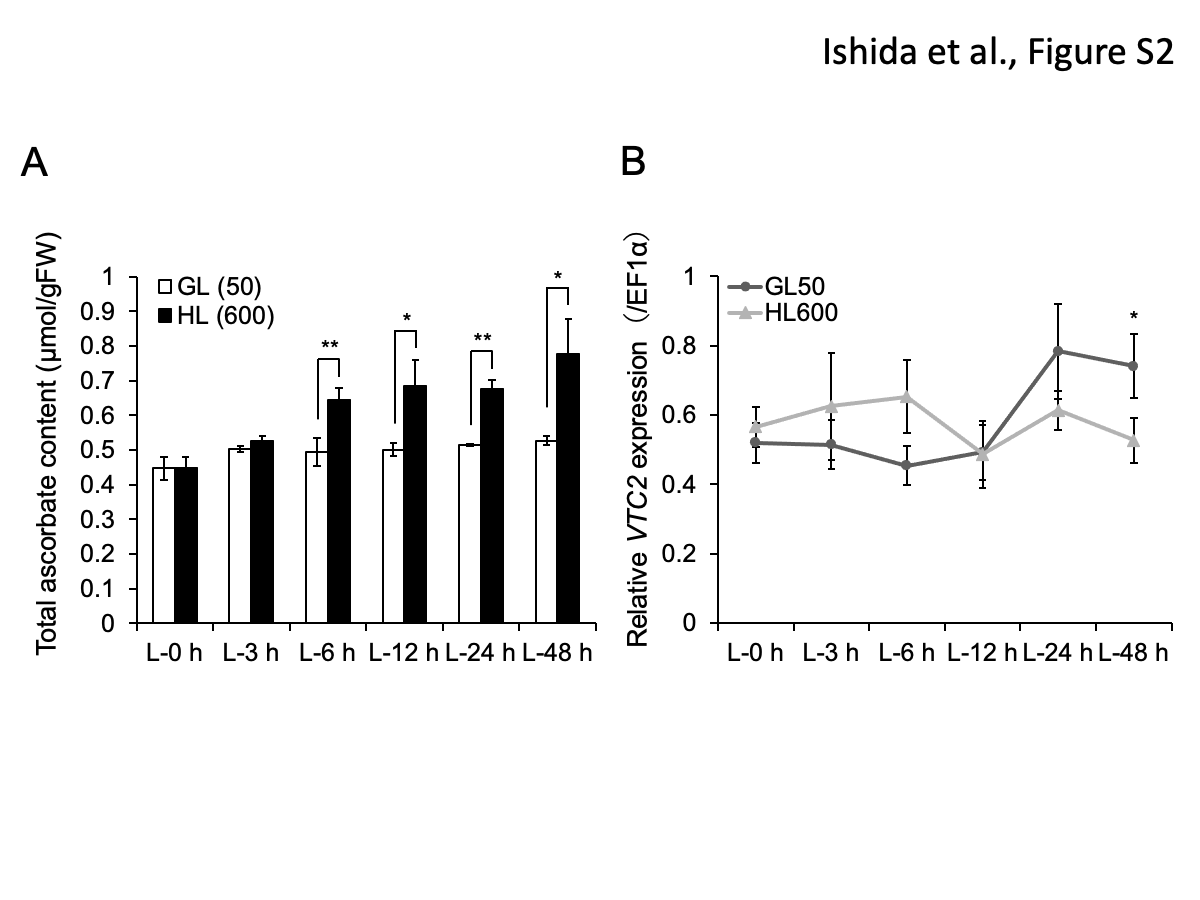

### Supplemental Figure S3

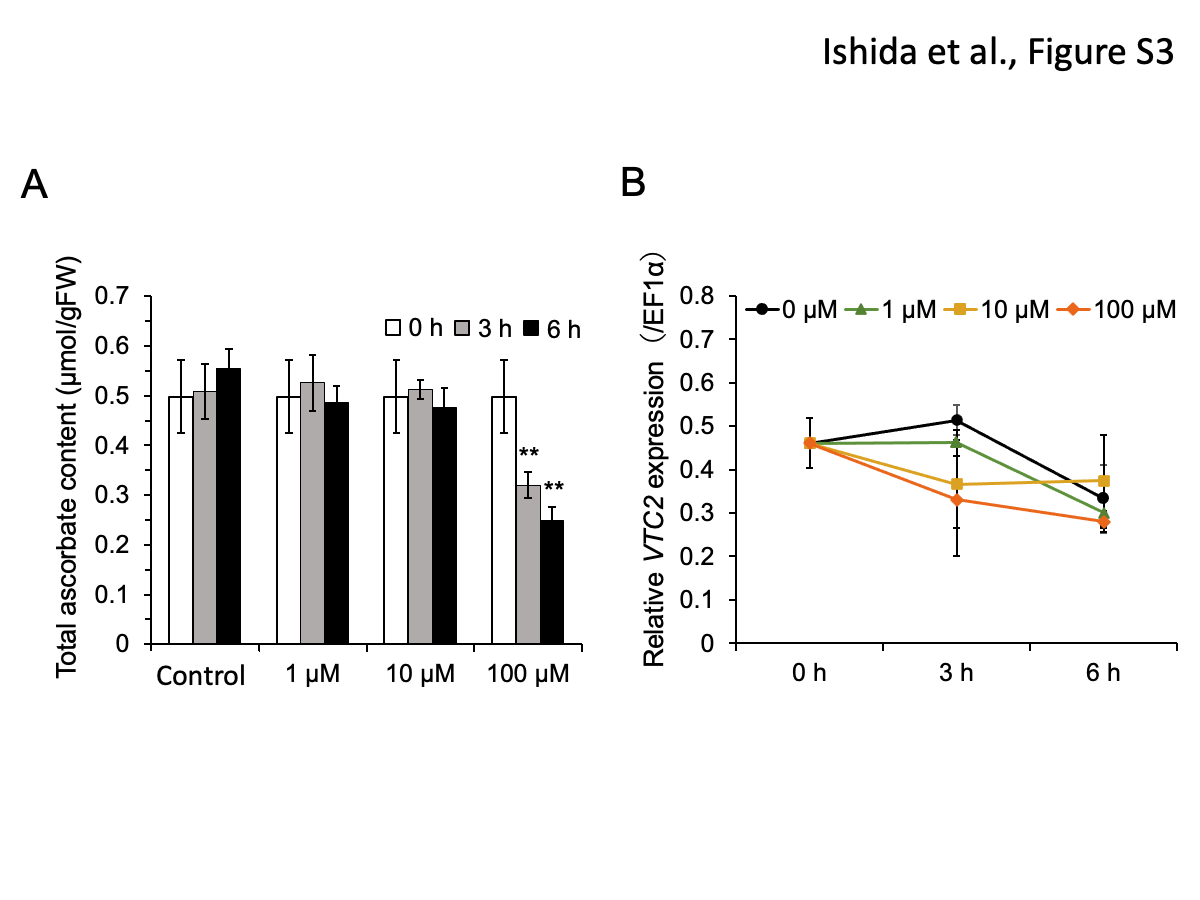
